# IsoformSwitchAnalyzeR v2: Analysis of Functional Isoform Changes in Long-read and Single-cell Sequencing Data

**DOI:** 10.64898/2025.12.08.693027

**Authors:** Chunxu Han, Jeroen Gilis, Elena Iriondo Delgado, Lieven Clement, Kristoffer Vitting-Seerup

## Abstract

Alternative splicing enables a single gene to produce a variety of mRNA transcripts, significantly enhancing protein diversity in higher eukaryotes. Isoform switching refers to the differential usage of a gene’s transcripts and occurs pervasively across physiological and pathological conditions. IsoformSwitchAnalyzeR was developed to identify these isoform switches and analyze their functional consequences. Advances in RNA-seq technology, including long-read and single-cell sequencing, along with state-of-the-art computational tools, enable unprecedented accuracy in isoform switch identification and its functional consequences, necessitating an update to IsoformSwitchAnalyzeR. Here we present IsoformSwitchAnalyzeR 2.0, with substantial improvements in the robustness of isoform switch detection, the incorporation of new functional annotation types, and interoperability with other bioinformatics tools. We showcase how IsoformSwitchAnalyzeR’s standard workflow is now well-suited for analysis of both long-read RNA-seq and single-cell data through two case studies. Specifically, we analyze long-read data from patients with Alzheimer’s Disease and single-cell data from patients with glioblastoma. In both case studies, we find important isoform switches with disease-relevant functional consequences, showcasing the power of IsoformSwitchAnalyzeR v2. Taken together, these findings highlight the versatility and robustness of IsoformSwitchAnalyzeR in handling advanced sequencing technologies, thereby broadening its applicability across diverse research contexts.

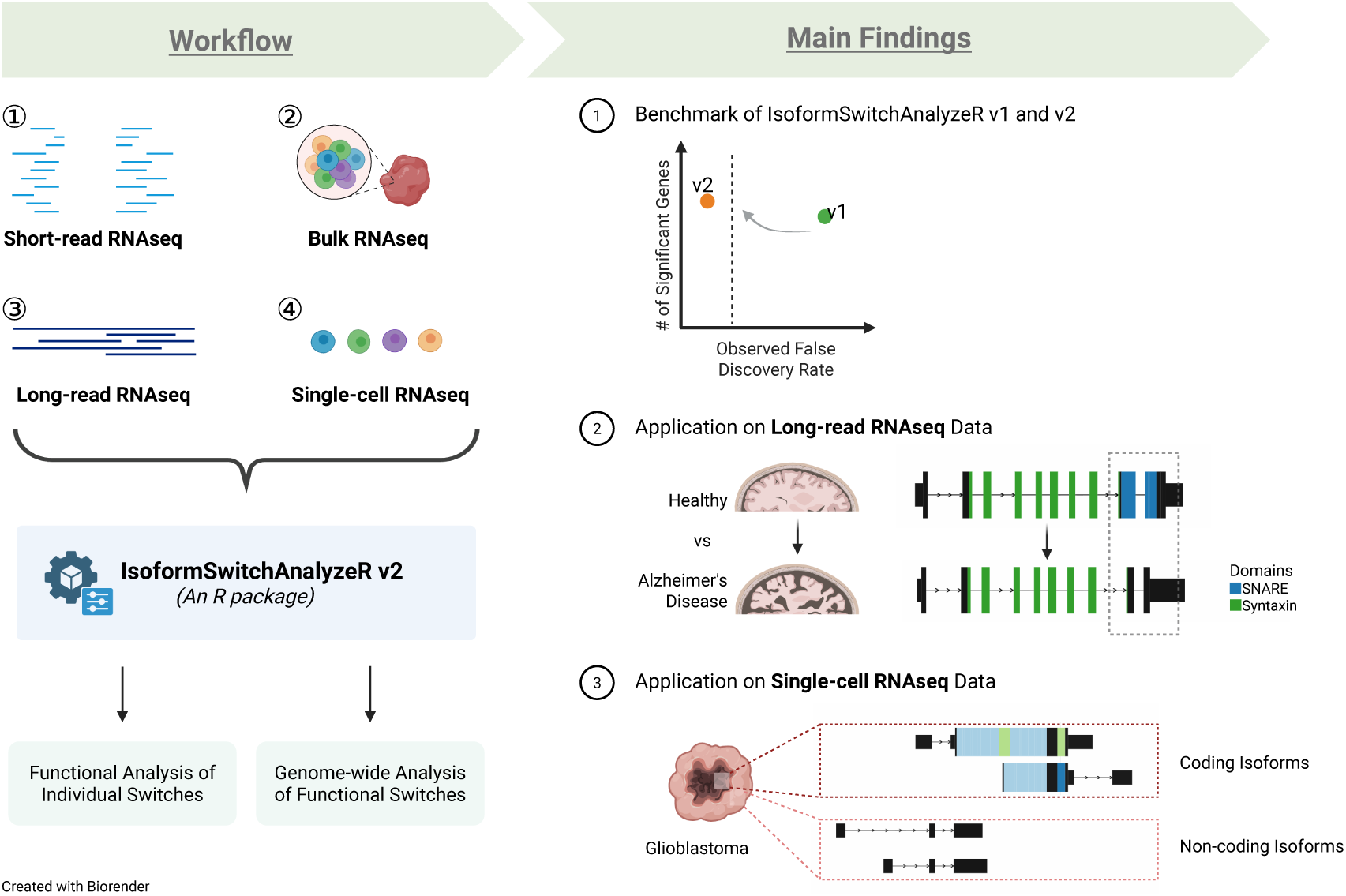

## Introduction

### Isoforms

Alternative splicing enables a single gene to produce multiple mRNAs, which are translated into distinct proteins. More than 95% of human genes with multiple exons undergo alternative splicing (1), and spliced isoforms of one gene may exhibit diverse structures, cellular localization, and biological functions (2). Differential usage of these isoforms, often referred to as "isoform switching, " is omnipresent across physiological and pathological conditions (3) and widely accepted as a “hallmark of cancer” contributing to many different aspects of cancer biology (4). As an example, Fujii *et al*. demonstrated a switch from CD45RA isoform to CD45RO isoform (with loss of exon 4) during T cell development and activation (5). Yanagisawa *et al*. demonstrated that among the isoforms of p120 catenin, only the full-length isoform, which includes both the central domain and the N-terminus, enhances binding stability and consequently inhibits RhoA activity, thereby facilitating increased cellular invasiveness (6). Barnkob *et al*. discovered that isoform switches in cell membrane proteins are significantly enriched, which may affect the targetable CAR therapy but also provide potential targets (7). Such findings highlight that analysis of isoform switches and their consequences can provide insights into cellular differentiation, disease progression, and drug development.

Despite hundreds of isoform switches with experimentally validated functional consequences existing in the literature (5–7), isoforms are still only rarely analyzed. As an example, our manual literature analysis found that in 2024 articles analyzing RNA-seq data, only 7% considered isoforms/splicing, but this trend generalizes to other research areas such as Alzheimer’s Disease (Figure 1A) and to the analysis of both common and rare genetic variation (8). To help close this gap, there is a clear need for bioinformatic tools that enable functional analysis of isoforms.

**Figure 1:**
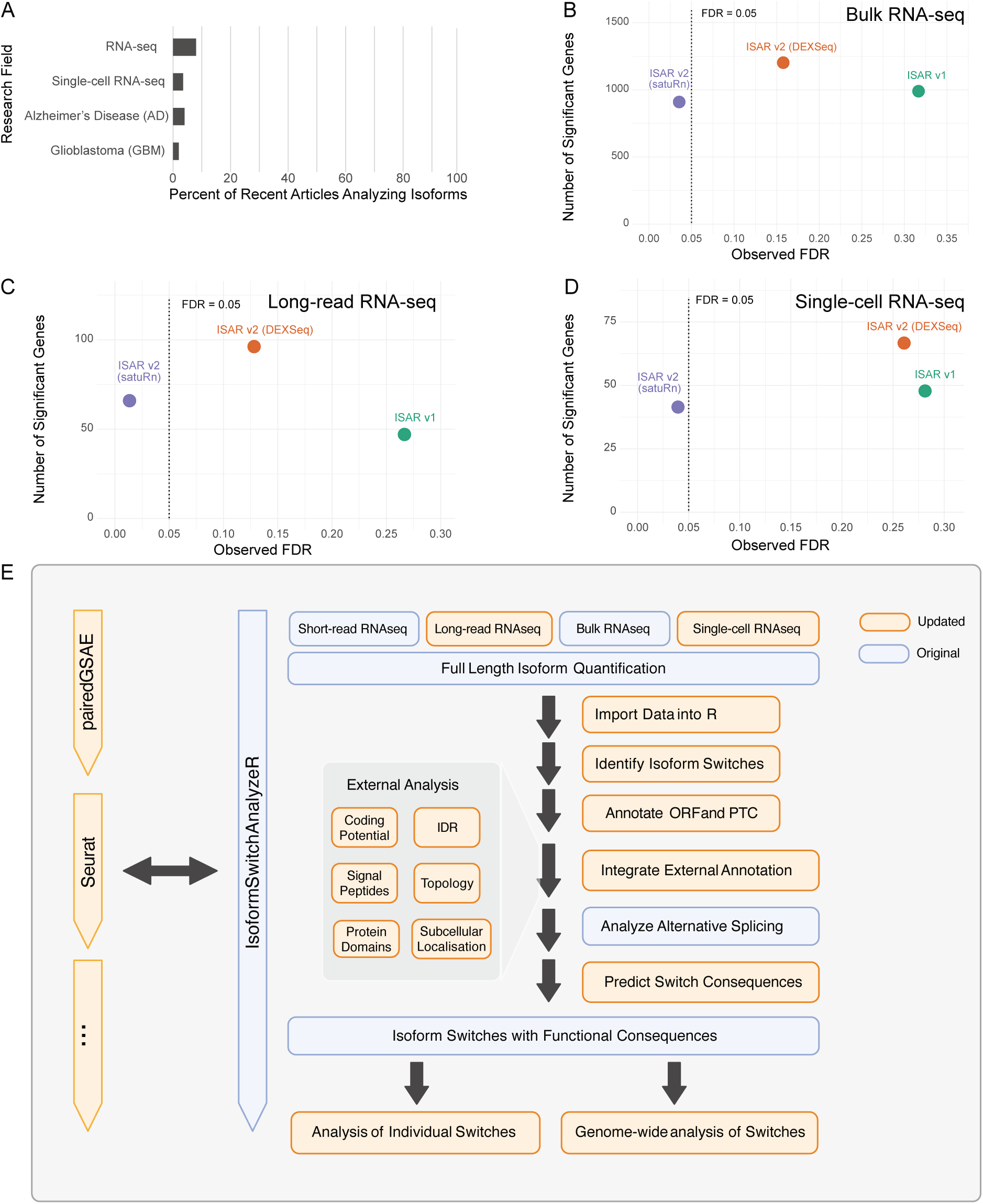
IsoformSwitchAnalyzeR 2.0 workflow, new functionality, and performance gain. **A)** Literature study of 300 articles showing how frequently isoforms were analyzed in articles about RNA-seq, single-cell RNA-seq, Alzheimer’s Disease, and Gliosblastoma. **B–D)** Comparison of the current and previous versions of IsoformSwitchAnalyzeR on bulk (B), long-read (C), and single-cell (D) RNA-seq, showing the number of significant genes identified as a function of the empirical (aka. observed) False Discovery Rate. The dashed line highlights the 5% nominal False Discovery Rate (aka. intended). **E)** The IsoformSwitchAnalyzeR pipeline now supports importing output from any tools that quantify full-length isoforms from different RNAseq data, including long-read and single-cell RNAseq. Following the identification of isoform switches, various tools for sequence annotation can be used to annotate the isoforms, thereby enabling the prediction of functional consequences. Once isoform switches with functional consequences are identified, two main types of analysis can be performed: analysis of individual switches and genome-wide analysis of switches. Orange boxes represent the new and updated features; original features are in blue.

### Analyzing the functional consequences of isoform switches

The first step towards analyzing the functional consequences of isoform switches is, naturally, the *identification* of such switches. Over the past decade, a large number have been created or repurposed towards this objective. This list is too long to recap exhaustively. Still, notable additions are DRIMseq (9), BANDITS (10), TSIS (11), Spycone (12), DEXSeq (13), EdgeR (14), and SatuRn (15), and these tools have been extensively reviewed for both bulk, long- and single-cell RNA-seq elsewhere (15–19).

To aid the biological interpretation of identified isoform switches, we, however, need bioinformatic tools that analyze the functional differences between isoforms used differentially across experimental conditions (e.g., healthy vs sick). This was the reason why, back in 2017, we created IsoformSwitchAnalyzeR (20). IsoformSwitchAnalyzeR works by identifying isoform switches, collecting functional annotations (e.g., protein domains), and systematically comparing the isoform annotations in the switch to identify differences (e.g., protein domain loss). We used this framework, e.g., to help explain the molecular function of isoform switches associated with worse survival rates in cancer patients, independent of cancer type (20). In 2019, we extended the IsoformSwitchAnalyzeR workflow with a module for genome-wide analysis. This analytical approach provides a genome-wide summary of all isoform switches and identifies statistically significant biases in functional consequences (e.g., more frequent domain loss than domain gain). Since then, two additional frameworks enabling the functional interpretation of isoform switches with predicted functional consequences have been published: 1) tappAS (21) in 2020, which includes an ingenious aggregation approach that potentially provides more power for differential feature analysis. 2) IsoTV (22) in 2021, which distinguishes itself by providing a streamlined processing pipeline.

Despite the increased availability of tools for analyzing isoform switches, IsoformSwitchAnalyzeR remains the most widely used. Accordingly, IsoformSwitchAnalyzeR has been applied across many research fields, revealing amongst others isoform changes in neurodegenerative diseases (23), cardiovascular diseases (24), virus infection (25), sepsis (26), venom evolution (27), various cancer types including oesophagal adenocarcinoma (28), oesophagal squamous cell carcinoma (29), as well as the cellular isoform dynamics during differentiation (30). These applications underscore its versatility and importance in understanding the isoform-level mechanisms underlying diverse biological conditions and diseases.

### Underlying developments

Since IsoformSwitchAnalyzeR was originally published, the field of transcriptomics has evolved considerably on both the technological and computational fronts. While bulk-RNA-seq remains a widely used and highly valuable method, innovations continue to revolutionize genome-wide investigations of alternative splicing and isoform characterization. Long-read sequencing enables the profiling of full-length RNA molecules without the need for error-prone assembly, thereby enhancing the detection of novel isoforms(31–34). The continued development of these technologies, along with software for quantifying and analyzing the data, means that long-read data is now routinely used to analyze differences between conditions (35). Additionally, using long-read data, even the most stringent analyses continue to identify thousands of new isoforms (36) – a trend likely to continue. While long-read sequencing enhances the accuracy and depth of isoform profiling, another technological breakthrough broadens isoform analysis: single-cell RNA sequencing (scRNA-seq). scRNA-seq measures the transcriptome of individual cells, revealing hidden cellular heterogeneity (37). And while the majority of current single-cell datasets cannot be used to quantify isoforms (38), some studies have examined isoforms using single-cell data, revealing a remarkable diversity (39, 40). Intriguingly, we are even starting to see long-read single-cell papers, and it is exciting to see how they advance our understanding of isoforms and their functional importance (41).

On the algorithm side, things have, if anything, moved even faster. Advanced computational tools such as Salmon (32) and kallisto (31) have become the primary quantification engines for bulk and single-cell RNA-seq. This also means that most researchers have isoform-level quantification of their data (although seldom using it (Figure 1A). We have seen the rise of new statistical methods for differential isoform usage (often called differential transcript usage, DTU) (10, 15, 42) and the repurposing of existing ones (13, 14, 43), leading to improved differential analysis. Lastly, tools for predicting isoform annotation have also seen tremendous progress, with, amongst others, improved tools for predicting protein domains (44), coding potential (45, 46), and signal peptides (47). Similarly, novel algorithms have been developed to predict various functional aspects of isoforms, including intrinsically disordered regions (IDRs) (48), protein topology (49), and subcellular localization (50), offering new and exciting ways to analyze isoforms.

Collectively, these technological and computational advancements improve the accuracy of isoform switch identification from both bulk-, long-, and single-cell RNA-seq, making it an ideal time to update IsoformSwitchAnalyzeR. Therefore, we present IsoformSwitchAnalyzeR 2.0, an updated version of IsoformSwitchAnalyzeR that now features additional functionality and an improved workflow for analyzing isoform switches with functional consequences and the underlying alternative splicing. We demonstrated the functionality of IsoformSwitchAnalyzeR 2.0 in two case studies: one on long-read sequencing of the Alzheimer’s disease cortex, followed by another on single-cell sequencing datasets of Glioblastomas. These studies showcase the features of IsoformSwitchAnalyzeR 2.0 and how the standard workflow can handle diverse and advanced sequencing technologies, thereby broadening its applicability in various research contexts.

## Methods

### Literature Survey

Multiple literature surveys were conducted to assess the prevalence of splicing-related analyses in RNA-seq studies, both overall and within specific disease contexts (e.g., Alzheimer’s disease, glioblastoma). Articles were retrieved from PubMed using the RISmed R package (v2.3.0). For each survey, inclusion criteria were defined to ensure relevance; these typically included (i) restricting publication dates to a defined time window (Jan–Jun 2024 for the RNA-seq and single-cell RNA-seq surveys; Jan 2022–Sep 2023 for the Alzheimer’s disease survey; Sep 2024–Sep 2025 for the Glioblastoma survey), (ii) article type limited to "Journal Article, " (iii) human studies only, and (iv) titles or abstracts containing terms indicative of RNA-seq (e.g., "RNA sequencing, " "bulk RNA sequencing" and "single-cell RNA sequencing") and, where applicable, disease-specific terms ("Alzheimer’s disease (AD)" or "Glioblastoma (GBM)").

From the retrieved articles, a random subset (n = 100 for the RNAseq survey and n = 50 for the disease survey) meeting the inclusion criteria was selected for manual review. Each article was assessed to confirm (a) it was a human study, (b) the full text was accessible, and (c) it used relevant RNA-seq data, and optionally, (d) it was relevant to the patients with certain diseases. Articles were then examined for evidence of splicing-related analyses by screening for terms such as “splicing, ” “isoform, ” or “transcript” in the Methods or other relevant sections. Studies were classified as performing splicing analysis if they included isoform- or transcript-level quantification, identification of alternative splicing events or changes, or other analyses beyond gene-level expression. The proportion of RNA-seq studies incorporating splicing analysis was then calculated as the number of such studies divided by the total number of reviewed articles (n = 100 or 50) for each survey.

### IsoformSwitchAnalyzeR

All analyses were performed with IsoformSwitchAnalyzeR 2.11.1 as available through Bioconductor (devel branch): https://bioconductor.org/packages/devel/bioc/html/IsoformSwitchAnalyzeR.html.

### Comparison of Statistical Methods for Differential Transcript Usage

To evaluate the performance of differential transcript usage methods across RNA-seq modalities, we simulated datasets from three sources: bulk RNA-seq from GTEx (51) expression profiles as detailed in (15); long-read RNA-seq from the GTEx long-read collection (https://gtexportal.org/home/downloads/adult-gtex/long_read_data); and single-cell RNA-seq from GSE106540/GSE106544, with transcript-level counts obtained from the ARCHS4 v2.2 database (52). For the single-cell modality, pseudo-bulk samples were obtained per donor by summing transcript counts across cells of the same homogeneous CD4⁺ TEMRA population.

For each modality, we generated 5 independent simulations by randomly sampling donors (3v3) into two artificial conditions. The lowly expressed transcripts were removed using edgeR’s (14) filterByExpr() function (default settings) as described in (15). Differential transcript usage was introduced in silico for 15% of multi-isoform genes by swapping transcript proportions within those genes, thereby mimicking realistic isoform-switching events. These transcript-level changes were encoded as a binary flag (txSwapped) in the metaInfo table, serving as the ground truth for evaluation at both transcript and gene levels.

We evaluated three DTU frameworks: IsoformSwitchAnalyzeR v2 (ISAR v2) using either DEXSeq or satuRn as well as the legacy ISAR v1 implementation. Each method was applied independently to all replicates using identical count matrices, transcript-gene mappings, and ground-truth DTUs. For gene-level analysis, we defined the gene-level DTU statistic (gene_switch_q_value) as the minimum isoform-level q-value among all transcripts of a gene, such that a gene is considered DTU if at least one of its isoforms is significant.

Performance was assessed using the following metrics: empirical True Positive Rate (TPR) and False Discovery Rate (FDR). TPR-FDR curves were generated using the iCOBRA R package (v1.30.0) (53), where each curve visualizes the sensitivity and precision trade-off of each method across false discovery thresholds. Additionally, we summarized performance using scatter plots of the number of significant true transcripts or genes (TP) versus observed FDR at q ≤ 0.05.

### Long-read RNA-seq data acquisition

The quantified long-read RNA-seq data were downloaded from Zenodo (https://zenodo.org/records/8180677) and contain datasets generated from the frontal cortex of 6 Alzheimer’s disease (AD) cases and 6 controls (54). We collected the transcript-level count matrix (counts_transcript.txt), in which isoforms were discovered and quantified by Bambu, based on version 107 of the Ensembl GRCh38 gene annotations (https://ftp.ensembl.org/pub/release-107/gtf/homo_sapiens/). We also retrieved the isoform nucleotide sequence (transcriptome.fa) and the genomic coordinates of the isoform exon structure (extended_annotations.gtf) from the transcriptomic_analysis_output folder as input to create the switchAnalyzeRlist object. The design matrix that indicates which samples belong to which conditions was established on the sample_metadata publicly available in Synapse (https://www.synapse.org/#!Synapse:syn52047893/files/). These data were used as input to a standard analysis workflow (see Figure 1) using IsoformSwitchAnalyzeR.

### Single-cell RNA-seq data acquisition and processing

The single-cell RNA-seq data were sourced from a study of surgical samples from 4 patients with confirmed primary glioblastoma (GSE84465) (55), in which, for each patient, two separate samples were collected: one originating from the tumor core and another from peripheral regions adjacent to the tumor core. The transcript-level count matrix was queried and extracted from the ARCHS4 (v 2.2) database based on GEO sample accessions (52). We utilized the cell type annotations and sample region information from the metadata provided by the original authors. We downloaded the GRCh38.107 annotation file, which was used to quantify the count matrix from Ensembl Archive (https://ftp.ensembl.org/pub/release-107/gtf/homo_sapiens/). To perform isoform switch analysis across different conditions for each cell type, we employed the AggregateExpression() function from the Seurat package to obtain pseudo bulk (56), which sums the gene counts of all cells from the same sample for each cell type. The data above were passed into IsoformSwitchAnalyzeR as the input.

### External Analysis

IsoformSwitchAnalyzeR extends its functionality by supporting the integration of output from external sequence analysis tools. Following the extraction of nucleotide sequences and the corresponding amino acid (AA) sequences of the ORFs, we leveraged a series of specialized bioinformatics tools to predict their functional consequences. We used Pfam’s pfam_scan.pl tool (v1.6) (57) to search the Pfam-A databases and predict protein domains. SignalP6 (v6.0) (47) and IUPred3 (v3) (48), with default settings, were employed to predict the presence of signal peptides and intrinsically disordered regions (IDRs), respectively. We applied DeepLoc2 (v2.0) (50) to predict protein subcellular localization using the High-throughput (Fast) model option. Lastly, to predict the topology of transmembrane proteins, we run DeepTMHMM (v1.0.24) (49) on the BioLib Cloud (https://dtu.biolib.com/DeepTMHMM) using the AA sequence file as input. Then we imported the output files from different tools and integrated the analysis into the switchAnalyzeRlist object via the built-in functionality in IsoformSwitchAnalyzeR via the analyze* function (where * is the name of the external tool, including PFAM, SignalP, and so on).

### Disease-relevance gene-set overlap analysis

To assess the disease relevance of the genes identified with functional isoform switches in our long-read Alzheimer’s disease (AD) and single-cell glioblastoma (GBM) analyses, we performed a gene-set overlap analysis for each disease using Gene Ontology Biological Process (GO:BP) gene sets retrieved from the Molecular Signatures Database (MSigDB) via the ‘msigdbr’ R package (v10.0.1). For each disease, well-established biological pathway categories in that disease were tested.

For Alzheimer’s disease: the following pathways are tested:

GOBP_NEURON_DEVELOPMENT, GOBP_NEURON_DIFFERENTIATION,
GOBP_AXON_DEVELOPMENT, GOBP_SYNAPTIC_SIGNALING,
GOBP_SYNAPSE_ORGANIZATION, GOBP_SYNAPSE_ASSEMBLY,
GOBP_NEUROTRANSMITTER_TRANSPORT,
GOBP_REGULATION_OF_SYNAPTIC_PLASTICITY,
GOBP_CALCIUM_ION_TRANSPORT,
GOBP_CALCIUM_ION_TRANSMEMBRANE_TRANSPORT,
GOBP_REGULATION_OF_CALCIUM_ION_TRANSPORT

For glioblastoma, the following pathways are tested:

GOBP_CELL_MIGRATION, GOBP_CELL_CYCLE,
GOBP_REGULATION_OF_CELL_POPULATION_PROLIFERATION,
GOBP_POSITIVE_REGULATION_OF_CELL_POPULATION_PROLIFERATION

For each pathway category, the union of all member genes was intersected with the set of DTU genes with predicted functional consequences.

### Visualizations

Figures 1E, 2A, 3A, and 3C were created in BioRender. The links to the figures with licenses are as follows: 1E: https://BioRender.com/4b42ofs, 2A: https://BioRender.com/k31r817, 3A: https://BioRender.com/731is5d, 3C: https://BioRender.com/qa3h2tk. All other figures were created with R using IsoformSwitchAnalyzeR’s built-in plotting functions.

**Fig. 2.**
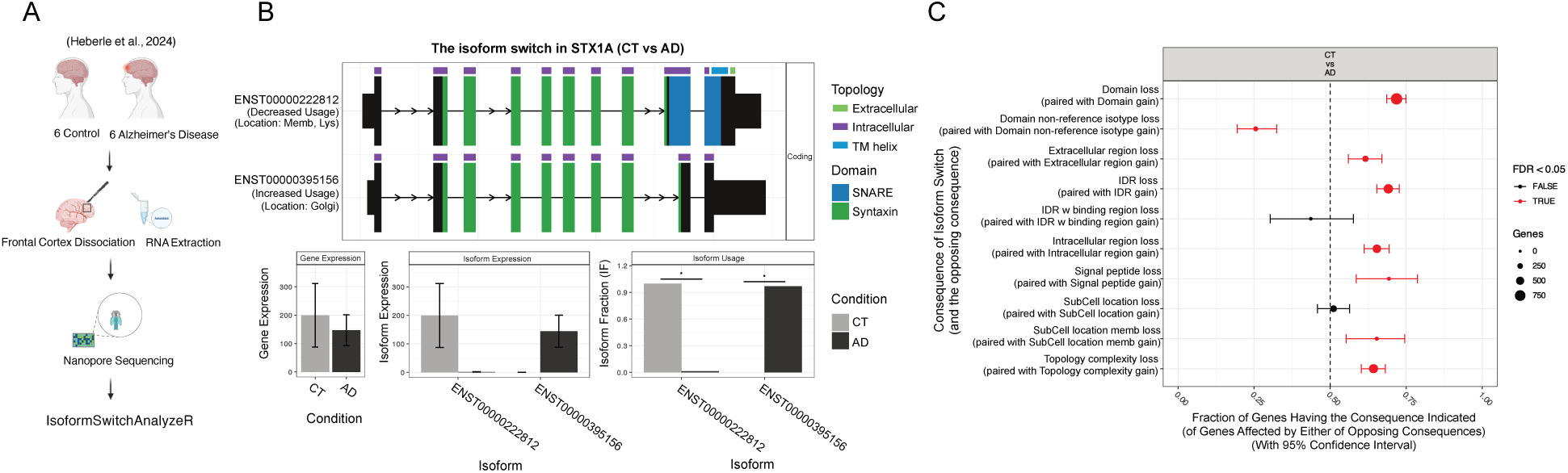
Isoform Switch Analysis on long read sequencing data on Alzheimer’s disease. **A)** Experiment design of long read sequencing data on the frontal cortex from Alzheimer’s disease (AD) and healthy controls. **B)** Isoform switch analysis of individual gene – STX1A. **C)** Gene-wide analysis of isoform switches with functional consequences. Here, we plot the fraction of genes (x-axis) with the indicated consequence (y-axis). The fraction is calculated as the fraction of genes with either of the paired opposite consequences. The FDR corrected p-values and confidence intervals are obtained from a binomial test. Panels B and C are shown exactly as created by IsoformSwitchAnalyzeR.

**Figure 3:**
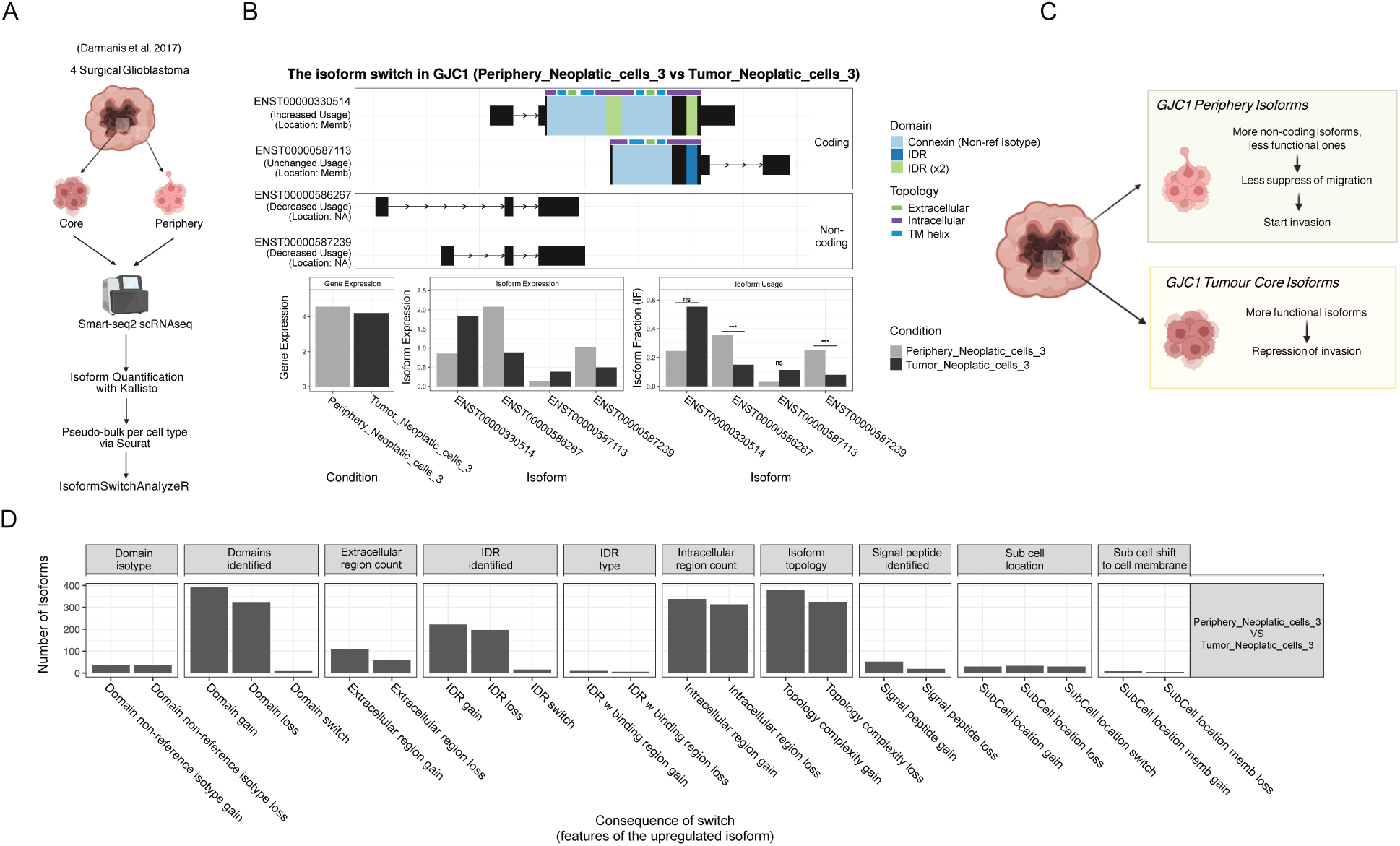
Single Cell Analysis of Isoform Switches by IsoformSwitchAnalyzeR v2. **A)** Overview of how the single-cell data reanalyzed here was generated and quantified. **B)** A switchPlot of the GJC1 gene created by IsoformSwitchAnalyzeR showing the annotated transcript structure (top), gene expression (bottom left), isoform expression (bottom middle), and isoform usage (bottom right). *** indicates FDR corrected p-value < 0.001. *ns* indicates “not significant”. **C)** Schematic illustration of our hypothesis for the function of GJC1 isoform in the tumor core and periphery. **D)** Genome-wide overview of the functional consequences in isoform switches between tumor core and tumor periphery created by IsoformSwitchAnalyzeR. Panels B and D are shown exactly as created by IsoformSwistchAnalyzeR.

### Quantifying the Prevalence of Structural Features Among Protein-Coding Genes

A SwitchAnalyzeRlist object containing all transcripts and sequences in GENCODE v41 was generated via IsoformSwitchAnalyzeR. Pfam’s pfam_scan.pl tool (v1.6) (57) and IUPred3 (v3) (48) were applied to predict the protein domains and intrinsically disordered regions (IDRs). The annotations were integrated into the isoformFeatures table. We calculated the percentage of protein-coding genes with at least one isoform containing a domain or an IDR by dividing the number of unique protein-coding genes with the respective annotation (domain_identified == "yes" or idr_identified == "yes") by the total number of protein-coding genes.

### Use of Generative AI

Generative AI was used to aid in the creation of this manuscript. Specifically, we used ChatGPT v4o and v4.5 as an editor and asked the model to improve the text of specific sections we had trouble formulating to our satisfaction. All text was carefully read to ensure ChatGPT added no meaning-changing edits or additions. ChatGPT and Perplexity.ai were also used, along with PubMed and Google Scholar, to search for relevant references. All references were read and verified before being added to the manuscript. Grammarly was used to spell-check and suggest improved phrasing, with all suggestions being carefully read and verified by the authors.

## Results

Here, we introduce IsoformSwitchAnalyzeR 2.0, an extended and improved version of the original tool (20). It is faster, produces more trustworthy results, and enables more insightful analysis of isoform switches with predicted functional consequences. Taken together, these updates make IsoformSwitchAnalyzeR the ideal tool for analyzing any transcriptomics dataset in which isoform quantification is available. Here, we first summarize the major changes in IsoformSwitchAnalyzeR v2 and subsequently demonstrate that it can analyze both long-read and single-cell RNA-seq data.

### Major updates in IsoformSwitchAnalyzeR version 2.0

Since the last major published release in 2019 (v 1.1.10), IsoformSwitchAnalyzeR has undergone a series of updates, ranging from minor bug fixes to major enhancements to the package’s capabilities and usage. Here, we will summarize only the major updates; a full list of more than 350 individual changes is included with the package. The major updates are also summarized in Figure 1E, which provides an overview of a typical IsoformSwitchAnalyzeR workflow, with parts that have undergone major updates highlighted in orange.

#### Improved robustness of analysis

After importing quantified isoforms and experimental metadata into IsoformSwitchAnalyzeR, the next step is the statistical identification of isoform switches (Figure 1E). This essential step of the IsoformSwitchAnalyzeR workflow has been improved in multiple ways, each contributing to the identification of trustworthy isoform switches. Firstly, we have recently shown that almost all RNA-seq datasets exhibit severe batch effects that, if not corrected for, render differential analysis of RNA-seq data unreliable (58). To accommodate this, we have updated IsoformSwitchAnalyzeR to handle both known (user-supplied) and unknown covariates (e.g., batch effects). Specifically, IsoformSwitchAnalyzeR now, in addition to user-supplied covariates, by default uses the SVA tool to infer surrogate variables to correct for unknown sources of variation, such as unknown batch effects (59). All covariates are included in the statistical analysis, ensuring that these unwanted effects do not affect downstream analyses.

Additionally, IsoformSwitchAnalyzeR uses limma’s removeBatchEffect (60) to remove the impact of unwanted covariates from the isoform expression data (TPM) that are used for both plotting and to calculate isoform switch effect size (dIF) (20). Finally, IsoformSwitchAnalyzeR was originally released with a statistical test we derived and implemented ourselves. But as the bioinformatic community became more aware of the challenge in differential transcript usage, new tools were developed (9, 10, 15), and existing tools were, after extensive benchmarking, repurposed (13). IsoformSwitchAnalyzeR has been continuously updated to align with best practices for DTU and now uses DEXSeq for smaller studies and satuRn for larger studies (where DEXSeq’s runtime is prohibitive), as well as for analysis of single-cell data (15, 19). These updates, combined with our continued focus on effect size cutoffs, result in the detection of more and more trustworthy isoform switches, as the False Discovery Rate (FDR) has decreased dramatically (Figure 1B-D, Supplementary Figure 1).

#### Extended support for functional consequence prediction

IsoformSwitchAnalyzeR v2 supports the analysis of 36 functional consequences (compared to 22 in v1, Supplementary Table 1). The consequences are largely enabled by incorporating external predictions, such as protein domains from Pfam (57) and signal peptides from SignalP (47), and then comparing the isoform in an isoform switch to identify annotation differences (e.g., domain loss). Supplementary Table 2 lists all external tool dependencies. While we have naturally maintained IsoformSwitchAnalyzeR to support the latest versions of these annotation tools (e.g., the update from SignalP4 (61) to SignalP5 (62) and SignalP6 (47)), we have also included many new annotations to improve the prediction of functional consequences, which we will briefly outline below.

Intrinsically disordered regions (IDR) are defined as regions that lack a stable 3D structure and, hence, are the exact opposite of protein domains. But, like protein domains, IDRs are extremely widespread and participate in almost all molecular functions (63). IDRs are particularly important for protein-protein interactions, cellular signaling, and liquid-liquid phase separation (64–66). To understand the molecular function of isoforms, it is therefore paramount to also understand IDRs; hence, support for IDR annotation has been built into IsoformSwitchAnalyzeR. Specifically, we support IDR annotations from NetSurfP-3 (67) and IUPred3 (48). NetSurfP-3 is supported because it has higher accuracy, but IUPred3 also allows predicting IDR, which facilitates interactions and thereby improves interpretability. Given the biological importance of IDR and the fact that 53% of protein-coding genes have at least one isoform with an IDR, we believe this is a major but often underappreciated source of isoform annotation.

Protein domains are often described as the building blocks of proteins and constitute an important, if not the most important, type of isoform annotation. This is partly because we find that 93% of protein-coding genes contain at least one isoform with a protein domain, and partly because the biological function of many protein domains is known (e.g., DNA-binding domain). This view of protein domains has, however, turned out to be a bit too simplistic, since 80% of human protein-coding genes use different subtypes of the same protein domain, termed domain isotypes (44). To accommodate this, IsoformSwitchAnalyzeR has been extended to support analysis of protein domain isotypes via pfamAnalyzeR (44), thereby revealing an extra dimension to protein domain analysis.

Another intriguing development in protein annotation is the ability to predict the subcellular localization of protein domains, thereby identifying isoforms that change their subcellular localization (50). Since this allows for unique hypotheses about the function of isoform switches, we have also incorporated support for sub-cellular predictions via DeepLoc2 (50)

Lastly, we recently showed that isoform switches in human cancers often change the topology of surface proteins by reducing the size of the extracellular part of the protein, which has important implications for cancer immunotherapy (7). To enable such analysis in other settings, IsoformSwitchAnalyzeR now supports topology annotation, which identifies which parts of a protein are intracellular, extracellular, or transmembrane, as well as the identification of transmembrane helices via DeepTMHMM (49). Importantly, this analysis also directly supports the analysis of changes in subcellular localization and, in our experience, often helps explain them (e.g., loss of a transmembrane helix explains changes in subcellular localization).

#### Interoperability with other bioinformatic tools

Growing evidence shows that isoform-level analysis provides an additional dimension to gene-level analysis of omics data (8, 58). These findings also highlight the need for new tools and databases that can systematically characterize isoform changes across biological contexts. To avoid the proliferation of isolated tools, we have established the isoformUniverse initiative, available as an R package through GitHub (https://github.com/IsoformAnalysisGroup/isoformUniverse). The isoformUniverse aims to provide a coordinated ecosystem of R packages for integrated, comprehensive, and multi-faceted isoform analysis across data modalities, biological contexts, and disease settings. To ensure that IsoformSwitchAnalyzeR v2 is part of this broader ecosystem rather than an analytical silo, we have integrated it into isoformUniverse. Although the initiative is still expanding, at the time of writing, it consists of IsoformSwitchAnalyzeR and PairedGSEA (58). PairedGSEA is an R package that makes a paired analysis of differential gene expression (DGE) (68) and differential isoform usage (13) and subsequently does Gene Set Enrichment Analysis (GSEA) (69) on both DGE and DTU results, making the result easier to interpret in a biological context. This global analysis of both DGE and DTU, along with the biological insights gained from the subsequent GSEA, complements the consequence-focused analysis performed by IsoformSwitchAnalyzeR extremely well. To allow users to utilize this synergy, we have updated IsoformSwitchAnalyzeR to both export data for analysis in pairedGSEA and import the results back. In other words, pairedGSEA and IsoformSwitchAnalyzeR are now fully interoperable and constitute a comprehensive analysis pipeline for transcriptomic data, but can naturally still be used independently, as is done in this article.

We have recently shown that isoform analysis, in addition to gene-level analysis, provides a second dimension to analyzing genomics and transcriptomics data (8, 58). This is especially true for transcriptomics data, where we found that gene expression and splicing mediate distinct biological signals (58). This result was obtained by another R package we created: PairedGSEA. PairedGSEA is an R package that makes a paired analysis of differential gene expression (DGE) (68) and differential transcript usage (7) and subsequently does Gene Set Enrichment Analysis (GSEA) (69) on both DGE and DTU results, making the result easier to interpret in a biological context. This global analysis of both DGE and DTU, along with the biological insights gained from the subsequent GSEA, complements the consequence-focused analysis performed by IsoformSwitchAnalyzeR extremely well. To allow users to utilize this synergy, we have updated IsoformSwitchAnalyzeR to both export data for analysis in pairedGSEA and import the results back. In other words, pairedGSEA and IsoformSwitchAnalyzeR are now fully interoperable and constitute a comprehensive analysis pipeline for transcriptomic data but can naturally still be used independently.

Taken together, these updates, along with many smaller ones, make IsoformSwitchAnalyzeR v2 a faster, more reliable tool that enables the discovery of isoform switches with many new types of functional consequences. These updates also make IsoformSwitchAnalyzeR v2 suitable for analyzing other types of transcriptomics. More specifically, IsoformSwitchAnalyzeR v2, as we show below, can now analyze both long-read and single-cell RNA-seq data, generating additional biological insights and hypotheses.

### Analysis of long-read RNA-seq data

Alzheimer’s disease (AD) is a leading cause of age-related cognitive decline and imposes a substantial burden on healthcare systems in developed countries. Its etiology is complex, involving widespread molecular alterations. Although numerous isoform-level changes have been observed in AD patients (70–72), isoforms remain markedly understudied in this context (considered in only ∼4% of recent articles, Figure 1A).

To showcase the utility of IsoformSwitchAnalyzeR v2 for analyzing long-read sequencing data, we re-analyzed Oxford Nanopore long-read sequencing data from the frontal cortex of 6 Alzheimer’s disease (AD) and 6 healthy samples (54) (Figure 2A). We followed a standard IsoformSwitchAnalyzeR v2 workflow with satuRn (15) to perform differential transcript usage analysis, identifying 2384 significant isoform switches across 1396 genes (FDR < 0.05). Continuing with the standard workflow, we annotated the isoforms involved in the identified switch with, amongst others, annotations of protein domains (57), signal peptides (47), Intrinsically Disordered Regions (IDR) (48), protein topology (49), and subcellular localization (50). We then used IsoformSwitchAnalyzeR v2 to assess which isoform switches had predicted functional consequences, identifying 2134 switches across 1208 genes. Notably, 172 (14.2%) of these genes are involved in known AD-relevant pathways, including neuron development (73), synaptic signaling (74), and calcium signaling (75), with the remaining genes representing potential novel AD-relevant candidates detectable only at the isoform level.

Among these genes, a particularly interesting switch was found in the STX1A gene. STX1A encodes the Syntaxin-1 (Stx1) protein that is part of the SNARE complex (76). The SNARE complex is central to neuronal communication through neurotransmitters, and its disruption has been associated with AD and neurodegeneration (77, 78). The SNARE complex works by facilitating the fusion of the synaptic vesicles, containing neurotransmitters, with the presynaptic terminal end of the neuron, thereby releasing the neurotransmitters that activate the next neuron (76, 79). Comparing AD to controls, we found no expression differences at the gene level (Fig 2B, bottom left) but a large isoform switch (FDR-corrected P-value = 0.015; Fig 2B, bottom row, middle and right subplots). The switch from ENST00000222812, used in healthy brains, to ENST00000395156, used in AD brains, results in the production of an isoform that lacks the SNARE domain that is essential for assembling the SNARE complex(80, 81) (Figure 2B top). Based on the other annotation we gathered, we also predict that the AD-specific isoform lacks the transmembrane helix that normally anchors a membrane protein to membranes. Accordingly, the predicted sub-cellular localization also switches from membrane-bound to Golgi (Figure 2B, top panel). Taken together, these observations support the hypothesis that the AD-specific isoform of Syntaxin-1 loses its function and therefore cannot participate in the SNARE complex, thereby inhibiting the release of synaptic vesicles and potentially contributing to AD pathogenesis.

This is, however, just one example of an isoform switch with a potential functional consequence. Summarizing across the functional consequences of all 1, 208 genes containing such switches, we find that the functional consequences of isoform switches are not random. There is a very clear and highly significant pattern in which isoforms lose functional features (e.g., domain loss, IDR loss, etc.) (Figure 2C). Part of this explanation is that the AD isoform, compared to its control counterpart, utilizes alternative transcription termination sites much more frequently (Supplementary Figure 2) and produces shorter isoforms.

Together, these results demonstrate how IsoformSwitchAnalyzeR v2 enables comprehensive analysis of both gene-specific and transcriptome-wide isoform usage changes from long-read RNA-seq data. The framework not only facilitates robust detection and annotation of functionally relevant isoform switches but also supports data-driven hypothesis generation regarding their potential roles in disease mechanisms—such as impaired synaptic function in Alzheimer’s disease.

### Analysis of single-cell RNA-seq Data

Glioblastoma is among the most lethal cancers, partly due to the critical location of the tumor within brain tissue, which limits the extent of surgical resection. As a result, infiltrating peripheral tumor cells are frequently left behind, leading to rapid recurrence within 6–9 months (82, 83). These residual cells are known to differ transcriptionally from tumor core cells (84). To add to these findings, we investigated isoform-level differences between core and peripheral glioblastoma cells using SMART-seq2-based single-cell RNA-seq data from four patients with paired core and peripheral tumor samples (55) (Figure 3A).

The single-cell isoform-level quantifications, performed with kallisto, were pseudo-bulked per cell type using Seurat, and the resulting count matrix was used as input to a standard IsoformSwitchAnalyzeR workflow using satuRn, as it is the best statistical engine for single-cell data (Figure 1D) (15, 19). We identified 1, 044 isoform switches with predicted functional consequences from 732 genes. Among these, 183 (25.0%) genes participate in cancer-relevant pathways, including cell migration (85) and cell proliferation (86), which are recognized hallmarks of glioblastoma biology, while the remaining genes represent potential novel candidates whose isoform-level dysregulation may contribute to the invasive phenotype of peripheral GBM cells. An intriguing example is in the GJC1 gene, which is associated with poor patient prognosis (87). GJC1 encodes a protein that is a component of gap junctions and is thereby involved in binding cells together. Overexpression of GJC1 has been experimentally shown to reduce migration and invasion (88) and to play distinct roles in glioblastoma cancer stem cells (89). Here, we find that GJC1 gene expression between tumor core and peripheral tumor cells is identical (Figure 3B, bottom left), but at the isoform level, there are substantial differences (FDR-corrected P-value < 0.001; Figure 3B, bottom row, mid and right panels). Specifically, we find that peripheral tumor cells have increased expression of a non-coding isoform and consequently have less of the protein-coding isoform (Figure 3B top). This leads to the hypothesis that the effective downregulation of the protein-coding GJC1 isoforms in peripheral tumor cells could contribute to the invasive phenotype of these tumor cells – a hypothesis that naturally needs to be experimentally tested.

While GJC1 represents a striking example of isoform switching that leads to loss of coding potential, it is far from unique. Across the dataset, we identified hundreds of genes exhibiting similar functionally relevant isoform changes, including loss of coding regions, domains, and regulatory features (Figure 3D). These widespread alterations underscore the limitations of gene-level analysis and highlight the critical need for isoform-resolved approaches, both for single-cell RNA-seq in general and for investigating glioblastomas.

## Discussion

Here, we summarize the substantial extensions of IsoformSwitchAnalyzeR v2 and showcase how these updates enable novel analyses of both long-read and single-cell sequencing data. We furthermore demonstrate how IsoformSwitchAnalyzeR identifies disease-relevant isoform switches in patients with Alzheimer’s Disease and Glioblastoma and how the informative plots produced by IsoformSwitchAnalyzeR enable us to make mechanistic predictions of the functional consequences of these isoform switches. IsoformSwitchAnalyzeR furthermore allows us to analyze changes in global patterns of both predicted functional outcomes and the underlying changes in alternative splicing, thereby providing a holistic picture of isoform-level changes. Notably, most of these insights are only possible with IsoformSwitchAnalyzeR v2 due to its support for novel functional consequences, such as subcellular localization, isoform topology, and intrinsically disordered regions.

These findings also demonstrate that bioinformatic tools/databases that enable researchers to make biologically relevant interpretations of isoform-level changes are essential. While pairedGSEA combined with IsoformSwitchAnalyzeR v2 represents a good baseline, it is clear there is still much more work to be done to enable widespread isoform-level analysis. This becomes even more evident given the continuous increase in the number of annotated high-confidence isoforms enabled by long-read technologies (90). This is illustrated by the most recent estimate from the ENCODE long-read RNA-seq collection, which is that ∼89.9% of human protein-coding genes utilize multiple expressed isoforms (91). We expect this number to increase further as more diverse tissues/cell-types are robustly profiled with long-read sequencing.

Notably, the isoform switches highlighted here show no changes in gene expression (Figures 2B and 3B), meaning these changes would not be detected by standard gene-level analysis. These examples align well with our recent findings from large-scale meta-analysis of both transcriptomics and genomics data (8, 58). In these studies, we find that among all genes significantly affected by gene-level changes, isoform-level changes, or both gene and isoform-level changes, ∼45-50% of the genes have isoform-level changes, and ∼25% only have isoform-level changes, meaning they are not detected by standard gene-level analysis. This pattern, along with our findings (Figure 3) and recent observations (15, 39), also suggests that isoform-level analysis of single-cell data holds great potential. In the grander scheme of things, this study, along with many others (8, 58), demonstrates the need to shift from our current gene-centric research paradigm to an isoform-centric one.

## Acknowledgment

We want to thank Lars Rønn Olsen for his insightful feedback on early versions of the manuscript. To all the users who took the time to report problems or ask questions: Thank you! Without all that feedback, the package would not be nearly as good as it is today.

## Authors’ contributions

KVS conceived the study. KVS and CH implemented all updates to IsoformSwitchAnalyzeR except for the implementation of satuRn, which JG and LC did. CH, EID, CH, and KVS maintained IsoformSwitchAnalyzeR. CH did all analyses and interpreted the results together with KVS and JG. CH and KVS wrote the manuscript. All authors read, commented on, and approved the final manuscript.

