## Supplementary tables and figures for "IsoformSwitchAnalyzeR v2: Analysis of Functional Isoform Changes in Long-read and Single-cell Sequencing Data"

##### Supplementary Table 1

Comparison of IsoformSwitchAnalyzeR (v1 and v2), tappAS, and IsoTV across functional consequences supported by IsoformSwitchAnalyzeR v2. ✓ *= supported; — = not supported.*

| Category | Consequence | ISAR v1 | ISAR v2 | tappAS | IsoTV |
|---|---|---|---|---|---|
| **Transcript Structure** | `tss` | ✓ | ✓ | ✓ | ✓ |
| | `tts` | ✓ | ✓ | ✓ | ✓ |
| | `last_exon` | ✓ | ✓ | ✓ | ✓ |

| | | | | | |
|---|---|---|---|---|---|
| | isoform_length | ✓ | ✓ | ✓ | ✓ |
| | exon_number | ✓ | ✓ | ✓ | ✓ |
| | intron_structure | ✓ | ✓ | ✓ | ✓ |
| | intron_retention | ✓ | ✓ | ✓ | ✓ |
| | isoform_class_code | ✓ | ✓ | — | — |
| **Sequence Similarity** | isoform_seq_similarity | ✓ | ✓ | — | — |
| | ORF_seq_similarity | ✓ | ✓ | — | — |
| | 5_utr_seq_similarity | ✓ | ✓ | — | — |
| | 3_utr_seq_similarity | ✓ | ✓ | — | — |
| **Coding Potential** | coding_potential | ✓ | ✓ | — | — |
| **Open Reading Frame** | ORF_genomic | ✓ | ✓ | ✓ | — |
| | ORF_length | ✓ | ✓ | ✓ | — |
| | 5_utr_length | ✓ | ✓ | ✓ | — |
| | 3_utr_length | ✓ | ✓ | ✓ | — |
| **NMD** | NMD_status | ✓ | ✓ | ✓ | — |
| **Protein Domains** | domains_identified | ✓ | ✓ | ✓ | ✓ |
| | genomic_domain_position | ✓ | ✓ | ✓ | ✓ |
| | domain_length | ✓ | ✓ | ✓ | ✓ |
| | domain_isotype | — | ✓ | — | — |
| **Signal Peptide** | signal_peptide_identified | ✓ | ✓ | ✓ | — |
| **Intrinsically Disordered Regions** | IDR_identified | — | ✓ | ✓ | ✓ |
| | IDR_length | — | ✓ | ✓ | ✓ |
| | IDR_type | — | ✓ | ✓ | ✓ |

| | | | | | |
|---|---|---|---|---|---|
| **Sub-cellular Localization** | sub_cell_location | — | ✓ | — | — |
| | sub_cell_shift_to_cell_membrane | — | ✓ | — | — |
| | sub_cell_shift_to_cytoplasm | — | ✓ | — | — |
| | sub_cell_shift_to_nucleus | — | ✓ | — | — |
| | sub_cell_shift_to_Extracellular | — | ✓ | — | — |
| **Protein Topology** | isoform_topology | — | ✓ | — | — |
| | extracellular_region_count | — | ✓ | — | — |
| | intracellular_region_count | — | ✓ | — | — |
| | extracellular_region_length | — | ✓ | — | — |
| | intracellular_region_length | — | ✓ | — | — |
| **Summary** | Total number of consequences supported | 22 | 36 | 19 | 13 |

##### Supplementary Table 2

Functional consequences supported by IsoformSwitchAnalyzeR v1 and v2. For each consequence, we indicate whether it was available in v1 or in v2, the external tool(s) required, and the R function used to integrate the results into the IsoformSwitchAnalyzeR workflow. ✓ *= supported; — = not supported.*

| Category | Consequence | ISAR v1 | ISAR v2 | External dependencies | Integration Function |
|---|---|---|---|---|---|
| **Transcript Structure** | tss | ✓ | ✓ | — | Built-in |
| | tts | ✓ | ✓ | — | Built-in |
| | last_exon | ✓ | ✓ | — | Built-in |
| | isoform_length | ✓ | ✓ | — | Built-in |
| | exon_number | ✓ | ✓ | — | Built-in |

| | | | | | |
|---|---|---|---|---|---|
| | intron_structure | ✓ | ✓ | — | Built-in |
| | intron_retention | ✓ | ✓ | — | Built-in |
| | isoform_class_code | ✓ | ✓ | — | Built-in |
| **Sequence Similarity** | isoform_seq_similarity | ✓ | ✓ | — | Built-in |
| | ORF_seq_similarity | ✓ | ✓ | — | Built-in |
| | 5_utr_seq_similarity | ✓ | ✓ | — | Built-in |
| | 3_utr_seq_similarity | ✓ | ✓ | — | Built-in |
| **Coding Potential** | coding_potential | ✓ | ✓ | CPAT or CPC2 | analyzeCPAT() or analyzeCPC2() |
| **Open Reading Frame** | ORF_genomic | ✓ | ✓ | — | analyzeORF() |
| | ORF_length | ✓ | ✓ | — | analyzeORF() |
| | 5_utr_length | ✓ | ✓ | — | analyzeORF() |
| | 3_utr_length | ✓ | ✓ | — | analyzeORF() |
| **NMD** | NMD_status | ✓ | ✓ | — | analyzeORF() |
| **Protein Domains** | domains_identified | ✓ | ✓ | Pfam | analyzePFAM() |
| | genomic_domain_position | ✓ | ✓ | Pfam | analyzePFAM() |
| | domain_length | ✓ | ✓ | Pfam | analyzePFAM() |
| | domain_isotype | — | ✓ | Pfam | analyzePFAM() |
| **Signal Peptide** | signal_peptide_identified | ✓ | ✓ | SignalP5/6 | analyzeSignalP() |
| **Intrinsically Disordered Regions** | IDR_identified | — | ✓ | IUPred2A/3 or NetSurfP-3 | analyzeIUPred3() or analyzeNetSurfP3() |
| | IDR_length | — | ✓ | IUPred2A/3 or NetSurfP-3 | analyzeIUPred3() or analyzeNetSurfP3() |
| | IDR_type | — | ✓ | IUPred2A/3 | analyzeIUPred3() |
| | sub_cell_location | — | ✓ | DeepLoc2 | analyzeDeepLoc2() |

| | | | | | |
|---|---|---|---|---|---|
| Sub-cellular Localization | sub_cell_shift_to_cell_membrane | — | ✓ | DeepLoc2 | analyzeDeepLoc2() |
| | sub_cell_shift_to_cytoplasm | — | ✓ | DeepLoc2 | analyzeDeepLoc2() |
| | sub_cell_shift_to_nucleus | — | ✓ | DeepLoc2 | analyzeDeepLoc2() |
| | sub_cell_shift_to_Extracellular | — | ✓ | DeepLoc2 | analyzeDeepLoc2() |
| Protein Topology | isoform_topology | — | ✓ | DeepTMHMM | analyzeDeepTMHMM() |
| | extracellular_region_count | — | ✓ | DeepTMHMM | analyzeDeepTMHMM() |
| | intracellular_region_count | — | ✓ | DeepTMHMM | analyzeDeepTMHMM() |
| | extracellular_region_length | — | ✓ | DeepTMHMM | analyzeDeepTMHMM() |
| | intracellular_region_length | — | ✓ | DeepTMHMM | analyzeDeepTMHMM() |

### Supplementary Figures

#### Figure S1

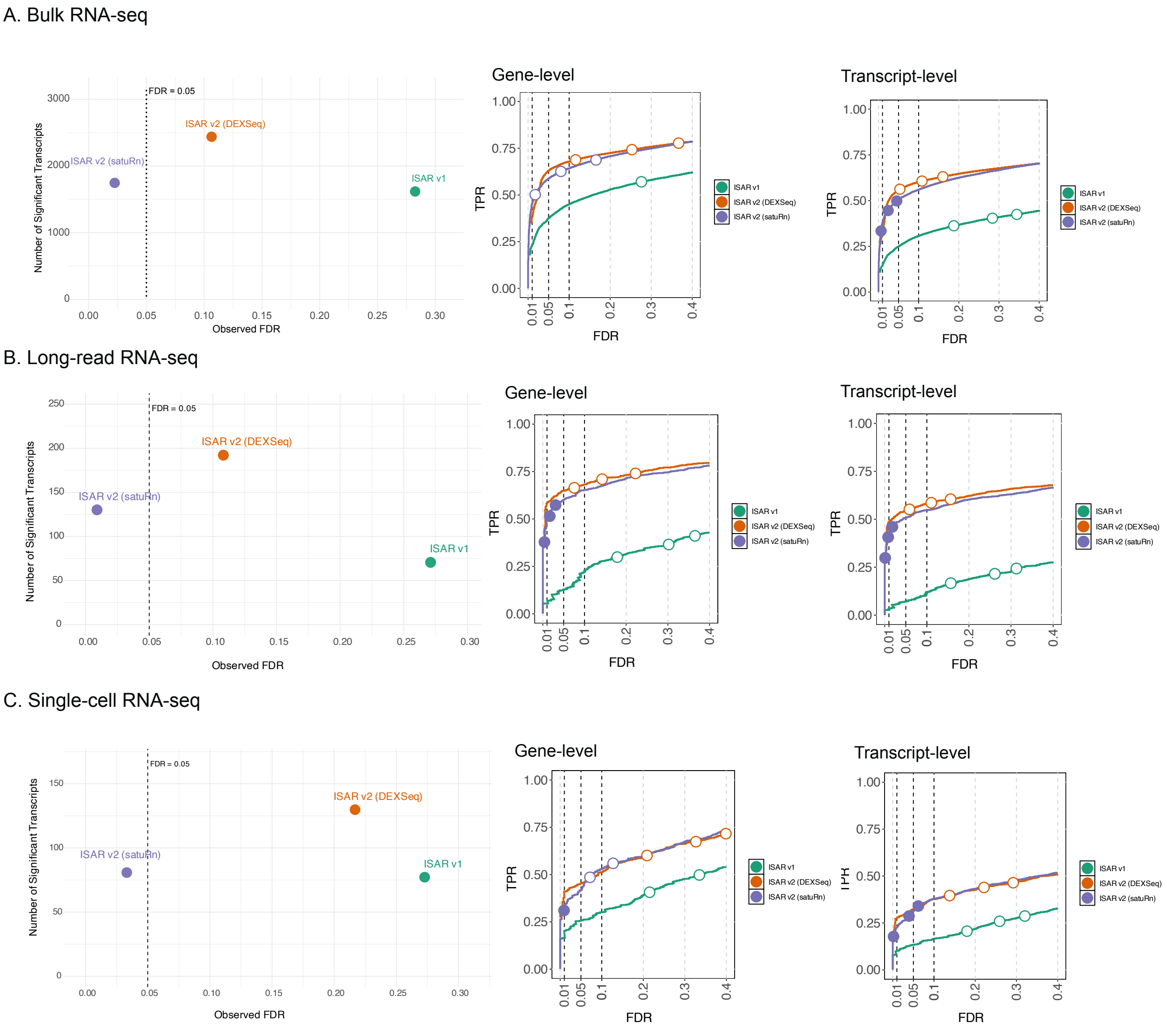

**Figure S1: Performance evaluation of ISAR v2 and ISAR v1 across three RNA-seq modalities.** Each row shows the performance of ISAR v1, ISAR v2 (DEXSeq), and ISAR v2 (satuRn) on simulated DTU built from **A)** bulk RNA-seq, **B)** long-read RNA-seq, and **C)** single-cell RNA-seq. For each modality, the left panel shows the number of significant transcripts (TP) identified at the 5% nominal FDR level as a function of the empirical False Discovery Rate; the dashed vertical line indicates the nominal FDR threshold. The middle and right panels show TPR–FDR curves at the gene and transcript levels, respectively, where the three circles on each curve indicate working points corresponding to nominal FDR levels of 1%, 5%, and 10%.

#### Figure S2

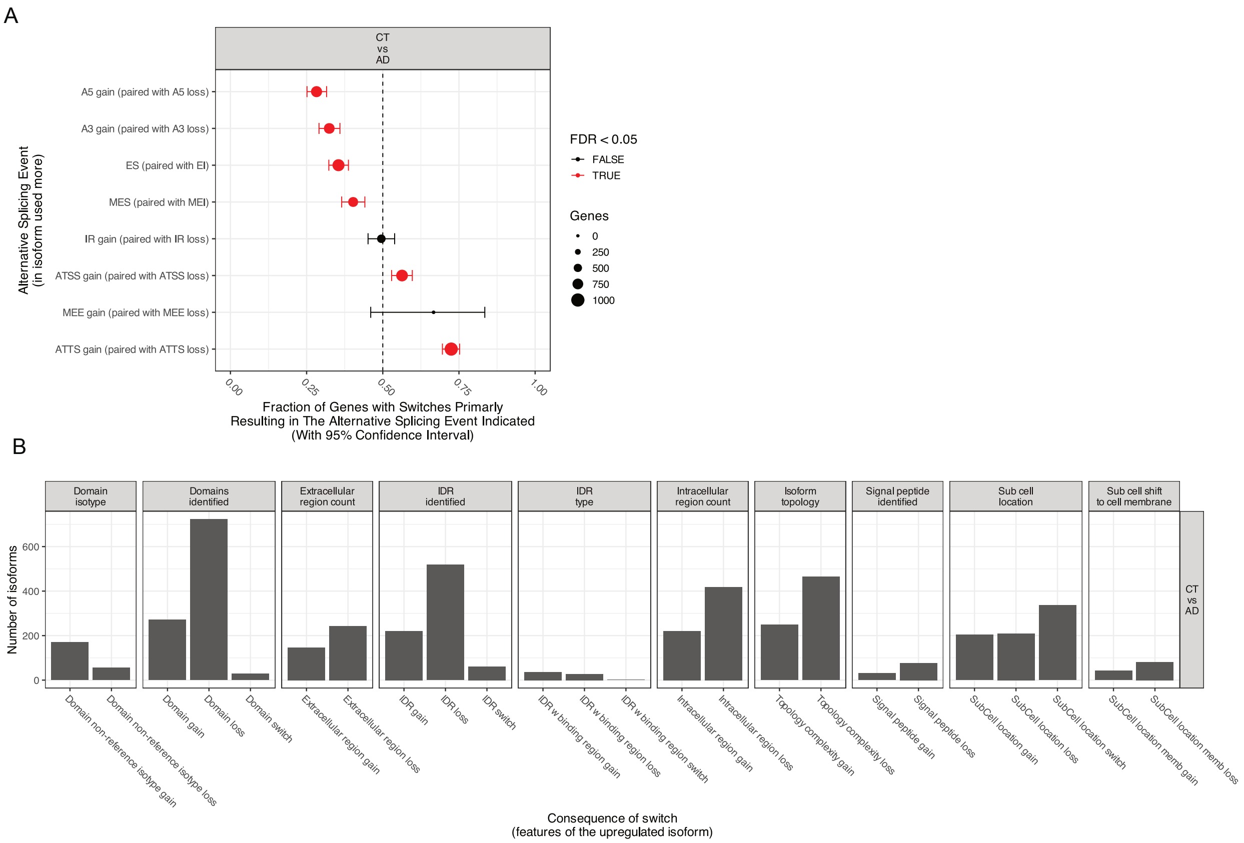

**Figure S2: Enrichment of splicing and consequences analysis of isoform switches by IsoformSwitchAnalyzeR v2 on long-read sequencing data on Alzheimer's disease. A)** The fraction (and 95% confidence interval) of isoform switches (x-axis) resulting in gain of a specific alternative splice event (indicated by y-axis) in the switch from control to AD. Alternative splicing events are abbreviated as follows: AS, alternative splicing; AI, alternative intron; ES, exon skipping; MEX, mutually exclusive exons; IR, intron retention; ATSS, alternative transcription start site; ATTS, alternative transcription termination site. **B)** Genome-wide overview of the functional consequences of isoform switches between AD and healthy controls. Bars represent the number of isoforms exhibiting specific gains or losses in features, including domain composition, intrinsically disordered regions (IDRs), signal peptides, subcellular localization, and membrane association.
